# ParSeek: Accurate cryo-EM particle picking with a deep learning model trained on synthetic data

**DOI:** 10.64898/2026.05.07.720949

**Authors:** Jiaqiang Qian, Yousheng Gong, Fuyan Liu, Yimiao Huang, Gaoxing Guo, Ya Zhu, Qiang Huang

## Abstract

Accurate particle picking from noisy cryo-EM micrographs is essential for high-resolution reconstruction. Current deep learning methods rely on manually annotated data, which is labor-intensive, subjective, and limits particle recall under low signal-to-noise ratio (SNR). Here we introduce ParSeek, an automated picker trained entirely on synthetic data without human annotation. Synthetic micrographs are generated by projecting known 3D structures into realistic background patches that reproduce experimental noise. On seven public cryo-EM datasets, ParSeek outperformed Topaz and CryoSegNet on four datasets, achieving the highest F1-score (up to 0.82) and reaching 0.63 on a challenging membrane protein dataset. Density maps from ParSeek-picked particles showed cross-correlation coefficients up to 0.995 with the reference and a minimal resolution difference of 0.1 Å. ParSeek also overcame severe orientation bias on an influenza dataset, yielding a reasonable reconstruction. Applied to three experimental datasets (an antibody–antigen complex and two GPCRs), ParSeek enabled reconstructions at 5.0 Å, 4.0 Å, and 2.8 Å, respectively. The 2.8 Å map resolved side-chain densities and ligand flexibility. This study establishes a fully synthetic-data-driven strategy that eliminates manual annotation for training cryo-EM deep-learning models, paving the way for automated, unbiased particle picking.

## Introduction

Cryo-electron microscopy (cryo-EM) has transformed structural biology by enabling high-resolution structure determination of macromolecules in their near-native state^1–3^. A typical cryo-EM experiment produces thousands of noisy micrographs. Accurate particle picking from these micrographs is a critical step, as its precision directly determines the quality of the final three-dimensional reconstruction^4^. To automate this process, deep learning has become the dominant paradigm. Existing methods typically adopt patch-based classification, semantic segmentation, or end-to-end object detection, significantly improving picking efficiency and generalization compared to traditional template matching. These models are typically trained to output particle probability maps or segmentation masks, from which particle coordinates are extracted for downstream processing^5^.

Representative deep learning-based pickers include DeepPicker^6^, Topaz^7^ and crYOLO^8^. These models are trained on manually annotated datasets. For example, DeepPicker used 1,084 micrographs from five datasets with expert-labeled particles and an equal number of negative samples. crYOLO required manual labeling of particles on approximately 900 micrographs across 45 datasets. Topaz was trained on about 300,000 positive particles from 3,000 micrographs across six datasets using a positive-unlabeled learning strategy. Manual annotation at this scale is labor-intensive and inherently subjective. Inter-operator variability is high, and many particles are missed under low signal-to-noise ratio (SNR), leading to low recall. Furthermore, manual labeling cannot scale to the growing volume of cryo-EM data, and each new macromolecular sample requires re-annotation. Consequently, the quality, completeness, and generalizability of training sets are fundamentally limited, constraining the performance of deep learning pickers.

To reduce manual labeling, many methods derive training annotations from single particle analysis (SPA) workflows or automated heuristics. For example, the CryoPPP^9^ dataset was constructed from SPA particle stacks and used to train CryoTransformer^10^ and CryoSegNet^11^. PIXER^12^ applied SPA to five EMPIAR datasets to generate its training set. However, SPA-derived annotations are biased because they come from 2D/3D classification, which preferentially selects particles that yield high-resolution reconstructions while discarding rare conformations. The classification process also requires expert intervention and is subjective. Other methods use automated pseudo-labeling, such as CASSPER (image-based metrics)^13^, U-Picker (A-LoG operator)^14^, or REPIC (consensus across pickers)^15^. Yet these heuristics often have low recall, require parameter tuning or seed labels, and cannot eliminate bias. Thus, neither approach overcomes the fundamental limitations of manual labeling: low recall, dataset-specific bias, and poor generalizability.

Given the limitations of manual and semi-automated annotations, synthetic data has been explored in cryo-EM. However, most efforts have focused on simulating imaging physics or generating realistic micrographs rather than specifically training particle pickers. Tools such as TEM-simulator^16^ and Parakeet^17^ generate synthetic micrographs by mimicking physical imaging processes. More recently, VirtualIce^18^ introduced experimentally acquired blank micrographs as realistic backgrounds. Yet these simulators were not designed to train a particle picker, and their outputs have not been validated for that purpose. Our previous work, PARSED^19^, provided an initial proof of concept by training a particle picker entirely on synthetic data, but it relied on unrealistic Gaussian noise backgrounds. Other methods have used synthetic data in more limited ways: BoxNet (Warp)^20^ incorporated a fraction of synthetic images to augment its training set, and U-Picker employed synthetic micrographs solely for benchmarking. Despite these efforts, existing synthetic data approaches are constrained by a disconnected workflow that separates data generation from model training. Rather than enabling an integrated pipeline where synthetic data and picking models are jointly optimized, existing methods either use synthetic data as a precomputed supplement, rely on unrealistic simulations, or do not train a picker at all. This fragmented design fundamentally fails to replace manual annotation.

Fully synthetic data offers distinct advantages: unlimited quantity, precise objective ground truth, and freedom from human bias. In computer vision^21^ and natural language processing^22^, models trained entirely on synthetic data have achieved performance comparable to real-data counterparts, enabling tasks such as object detection and semantic segmentation. In cryo-EM, synthetic data has similarly enabled deep learning models for other tasks, such as pose estimation (CryoFastAR)^23^ and density map generation (struc2mapGAN)^24^. This raises a natural question: can a similar fully synthetic approach enable high-performance particle picking in cryo-EM without any manual annotation? Here we present ParSeek, a particle picker trained exclusively on synthetic data. Our pipeline generates realistic micrographs by projecting known 3D structures into experimentally acquired background images and integrates a self-supervised denoising module with a segmentation network. ParSeek eliminates the need for manually labeled micrographs while achieving competitive performance on public benchmarks and challenging experimental datasets.

## Results

### Pipeline of ParSeek

The ParSeek pipeline consists of three stages (Fig. 1). First, to enhance raw micrographs and improve particle visibility without external training data, we applied a self-supervised denoising module. This blind-spot-based network suppresses position-independent noise by reconstructing pixels from local context (Figs. 1a and Supplementary Fig. 4c).

**Fig. 1.**
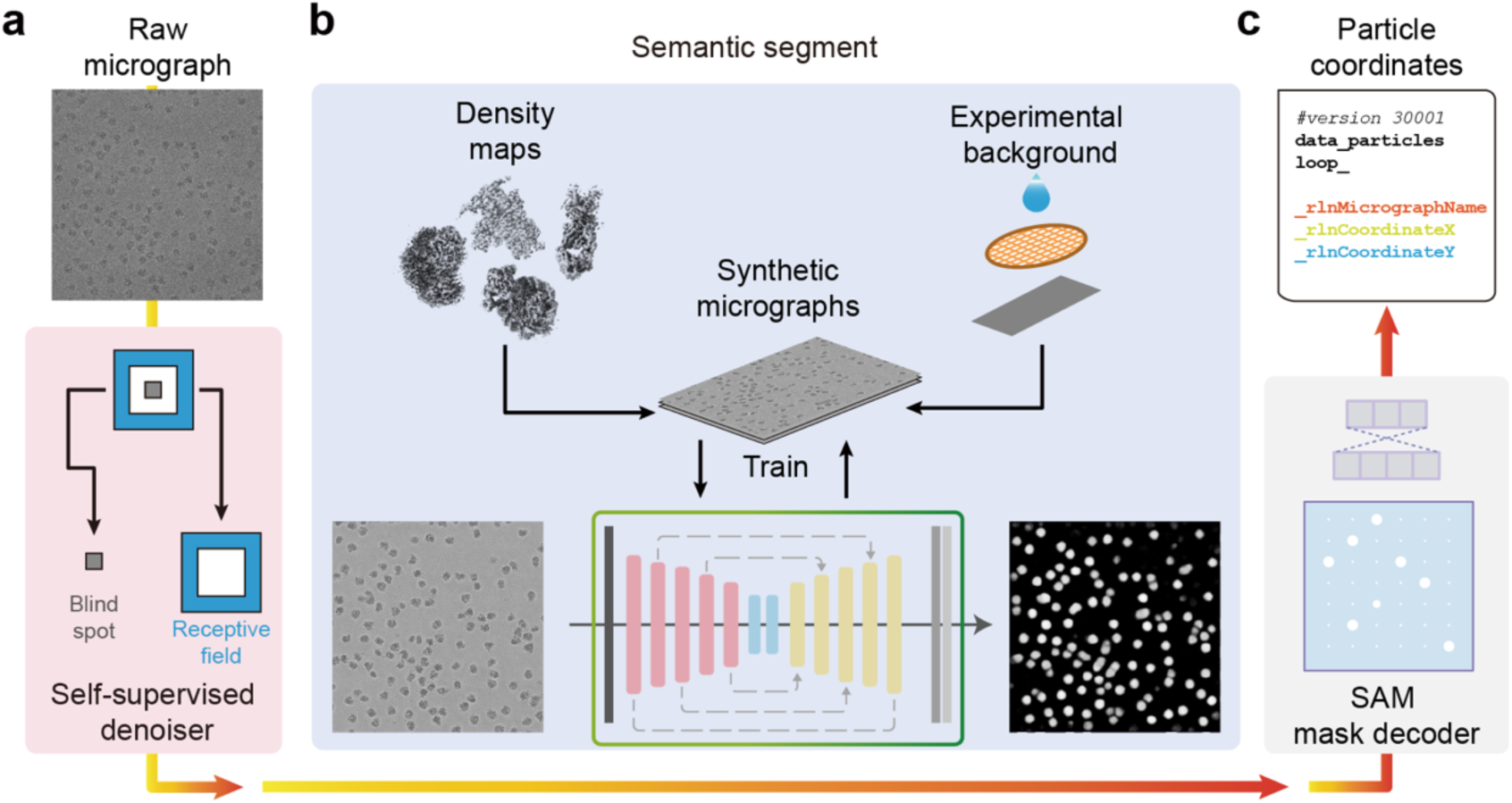
| Pipeline for self-supervised denoising and particle picking in ParSeek. **a**, Self-supervised denoising of a motion-corrected micrograph using a blind-spot neural network, depicting its input and output. **b**, Semantic segmentation of the processed image by a U-Net model trained on synthetic micrographs, outputting a binary mask of potential particles. **c**, Particle localization where a Segment Anything Model (SAM) converts the mask into precise coordinates for export in STAR format.

Second, ParSeek uses a U-Net-based semantic segmentation network for pixel-wise particle identification. To eliminate the need for manual annotation, the network was trained on a synthetic dataset generated by projecting diverse structural densities onto experimentally derived background micrographs, capturing variations in particle morphology and noise characteristics. The resulting model produces well-localized, circular binary masks that reliably segment particles even under challenging conditions (Fig. 1b).

Finally, particle coordinates are extracted using an adapted Segment Anything Model (SAM) framework^25^. This module divides each segmentation mask into a uniform grid, generates bounding boxes for candidate particles, and outputs standardized STAR files for direct use in RELION^4^ or cryoSPARC^26^ (Fig. 1c).

### Construction of synthetic micrographs

To overcome the fundamental bottleneck of manual annotation, we constructed a large-scale synthetic cryo-EM dataset for training semantic segmentation networks. Each synthetic micrograph combines two elements: experimentally derived backgrounds and computationally generated particle images (Fig. 2a). Background micrographs were sourced from buffer-only experimental data (EMPIAR-12287)^18^ and visually inspected to exclude regions contaminated with crystalline ice or carbon film edges (Supplementary Fig. S1). Particle images were generated from a diverse library of 22 macromolecular density maps (Supplementary Table S1). Density maps exhibiting significant signal loss in regions were replaced prior to projection (Supplementary Fig. S2).

**Fig. 2.**
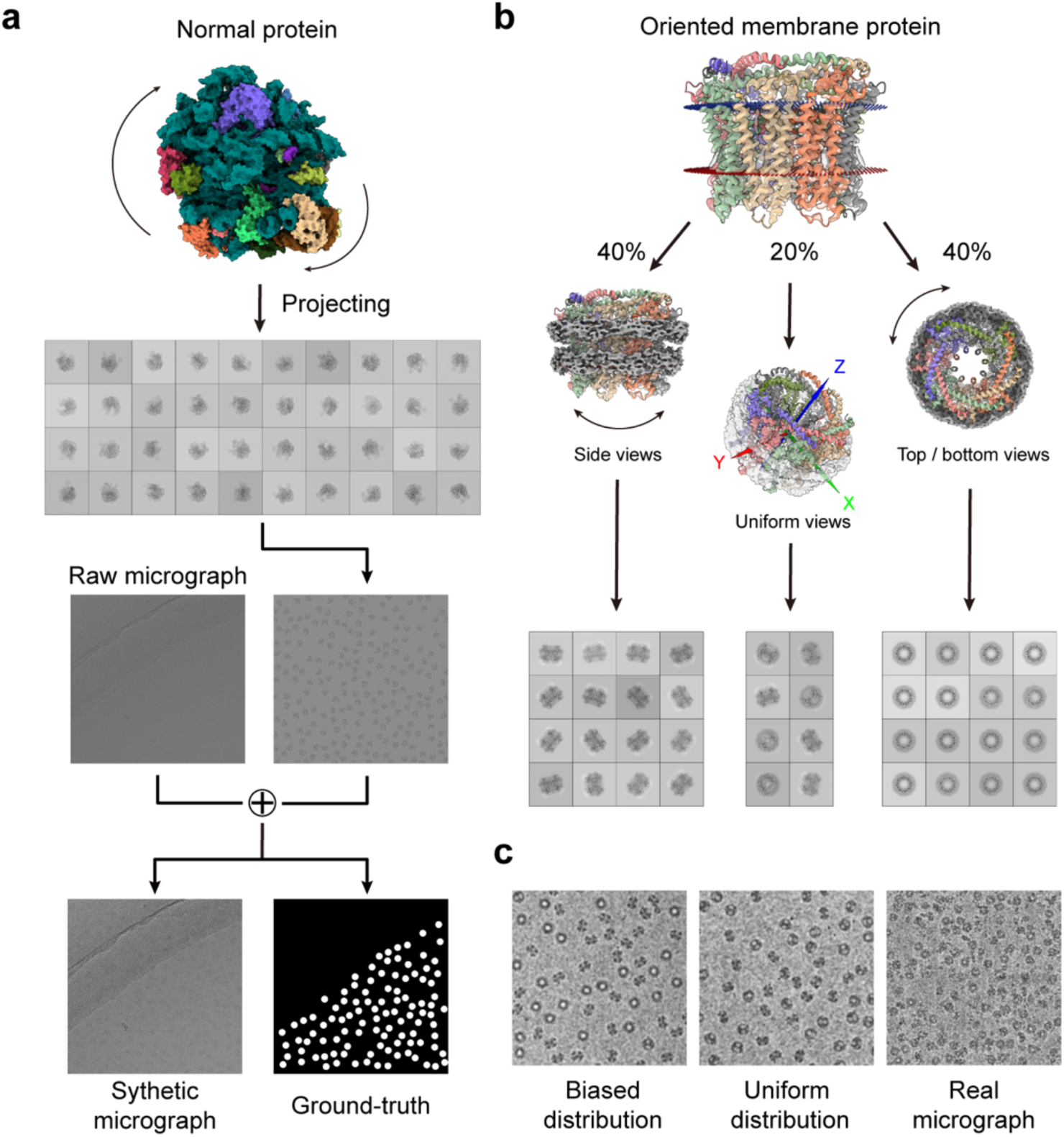
| Design and generation of synthetic micrographs. **a**, Pipeline for generating a synthetic micrograph and its ground-truth mask by integrating simulated particle images with experimental backgrounds. **b**, Membrane-protein-biased angular sampling strategy used for projection generation. **c**, Comparison of simulated particles from EMD-0919 under the biased (left) and uniform random (middle) sampling, alongside a real micrograph from EMPIAR-10444 (right).

In cryo-EM experiments, particle orientations are often biased due to interactions with the air–water interface, a phenomenon particularly pronounced for membrane proteins solubilized in detergent micelles, which frequently adopt preferred views (e.g., top and side orientations)^27^. To account for this, we applied a protein-type–aware angular sampling strategy: non-membrane proteins were sampled uniformly across all orientations to preserve view diversity (Fig. 2a), while membrane proteins were subjected to biased sampling that enriches top and side views commonly observed in experimental 2D class averages (Figs. 2b, c). This approach ensures full angular coverage for all targets while enhancing the realism of membrane protein views.

In the final assembly, these particle images were embedded into background micrographs at random locations within uncontaminated regions. For each synthetic micrograph, a corresponding ground-truth segmentation mask was generated (Fig. 2a), with valid particles represented as white circles on a black background. The mask format is directly compatibility with the U-Net training pipeline. The resulting synthetic dataset contains 6,600 unique micrograph–mask pairs and provides a fully annotation-free resource for particle picker training.

### Effects of synthetic data components

To evaluate the robustness of ParSeek under common cryo-EM conditions, we analyzed four representative micrographs from EMPIAR-10028 using two ablation models.

First, we assessed the impact of realistic background modeling by comparing ParSeek to a control model trained with Gaussian noise instead of experimental buffer backgrounds (denoted as ParSeek-Noise). While both models detected genuine particles in clean regions, ParSeek-Noise produced undersized masks and generated false positives by misclassifying noise features (Fig. 3a). This behavior demonstrates that without realistic background features, the model fails to generalize to experimental micrographs, highlighting the importance of experimental backgrounds for artifact suppression.

**Fig. 3.**
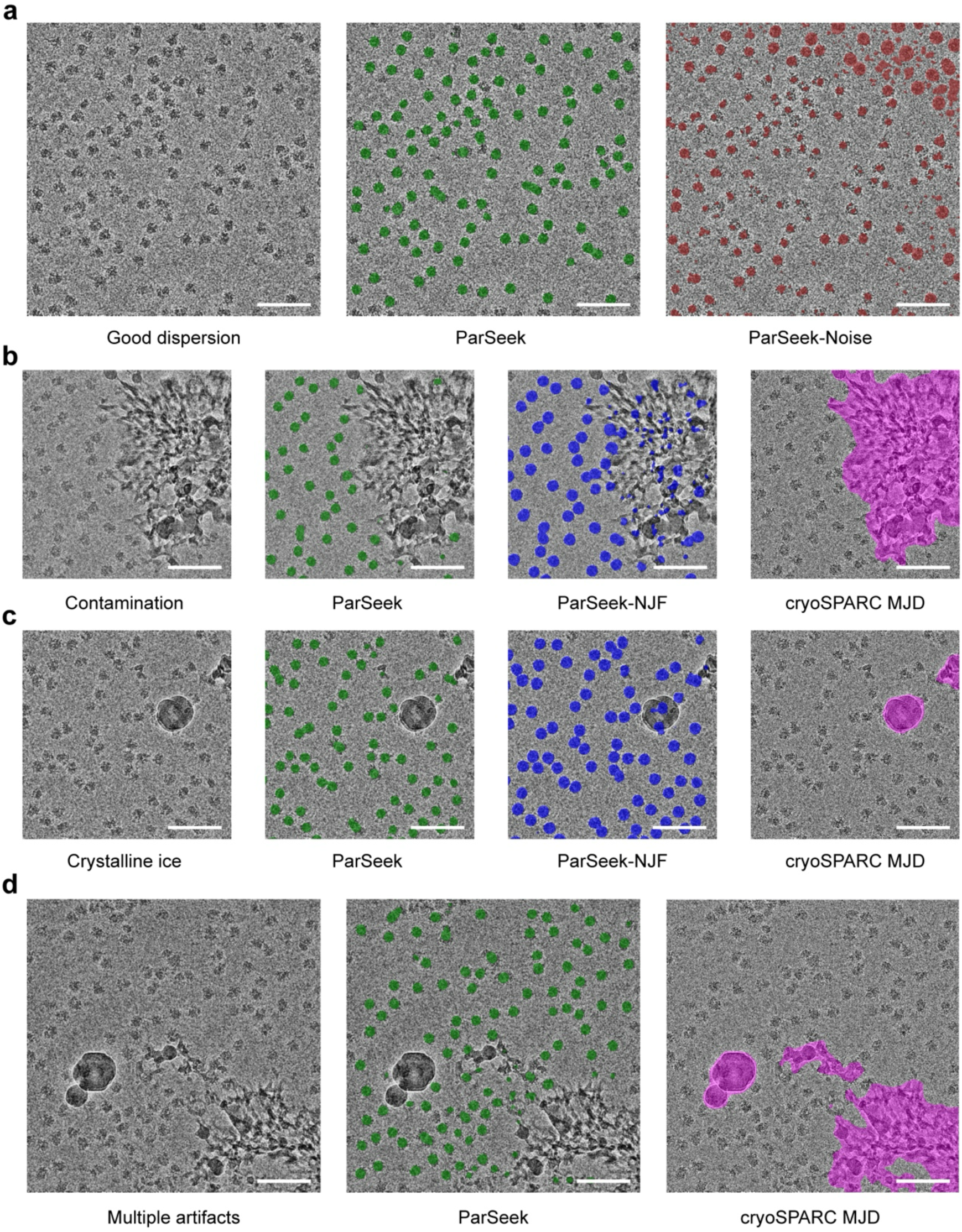
| Evaluation of ParSeek and ablated models on real, representative micrographs. **a**, A well-dispersed micrograph. From left to right: original image; ParSeek segmentation (green); ParSeek-Noise (trained on Gaussian noise background, brown). **b**, A micrograph with ice contamination. From left to right: original; ParSeek (green); ParSeek-NJF (trained without junk filtering, blue); cryoSPARC junk reference (magenta). **c**, A micrograph containing crystalline ice. From left to right: original; ParSeek (green); ParSeek-NJF (blue); cryoSPARC reference (magenta). **d**, A micrograph with hybrid pollution. From left to right: original; ParSeek (green); cryoSPARC reference (magenta). Scale bars, 100 nm.

Next, we evaluated the role of excluding contaminated regions during training by comparing ParSeek to a model trained without filtering out junk areas (denoted as ParSeek-NJF). ParSeek accurately restricted particle segmentation to uncontaminated regions, consistent with the junk mask generated by cryoSPARC Junk Detector (Figs. 3b, c). In contrast, ParSeek-NJF produced extensive false positives on both carbon contamination and crystalline ice. This result shows that training on unfiltered data causes the model to misinterpret artifacts as particles, confirming that explicit exclusion of contaminated regions is essential for learning genuine particle features. The full ParSeek model further demonstrated robustness on a micrograph with mixed contamination, correctly identifying valid particles while excluding defective areas (Fig. 3d).

Together, these comparisons demonstrate that the robust performance of ParSeek originates from two key design principles: (1) integrating real experimental micrograph backgrounds and, (2) excluding contaminated regions during synthetic data generation.

### Performance of ParSeek on benchmark datasets

We next compared ParSeek with Topaz and CryoSegNet, two established particle pickers trained on real experimental annotations, using the seven test sets from the CryoPPP benchmark (Supplementary Table S2). Topaz is widely used and integrated into RELION and cryoSPARC, and CryoSegNet was rigorously evaluated on the CryoPPP benchmark.

ParSeek achieved particle-picking accuracy comparable to state-of-the-art tools (Table 1), with representative detection outputs shown in Supplementary Fig. S4c. It obtained the highest F1-scores on four datasets (EMPIAR-10028, 10081, 10093, and 11056), while Topaz performed best on the remaining two (EMPIAR-10017 and 10345) and held a marginal lead (∼1%) on EMPIAR-10532. On datasets with higher overall F1-scores (>0.7, indicating fewer challenging conditions), the performance gap between ParSeek and Topaz was narrow. Conversely, on more challenging datasets with lower F1-scores, the two methods each outperformed the other on different subsets, demonstrating comparable robustness under difficult conditions.

**Table 1.** | Performance comparison of ParSeek, Topaz and CryoSegNet.

| EMPIAR<br>ID | ParSeek |  |  | Topaz |  |  | CryoSegNet |  |  |
| --- | --- | --- | --- | --- | --- | --- | --- | --- | --- |
|  | Precision | Recall | F1 score | Precision | Recall | F1 score | Precision | Recall | F1 score |
| 10017 | 0.73 | 0.72 | 0.73 | 0.77 | 0.75 | **0.76** | 0.72 | 0.17 | 0.28 |
| 10028 | 0.73 | 0.96 | **0.82** | 0.65 | 0.97 | 0.78 | 0.78 | 0.82 | 0.80 |
| 10081 | 0.66 | 0.87 | **0.75** | 0.60 | 0.89 | 0.72 | 0.67 | 0.79 | 0.73 |
| 10093 | 0.34 | 0.68 | **0.46** | 0.35 | 0.36 | 0.36 | 0.34 | 0.29 | 0.31 |
| 10345 | 0.27 | 0.75 | 0.40 | 0.51 | 0.86 | **0.64** | 0.40 | 0.56 | 0.47 |
| 10532 | 0.33 | 0.60 | 0.43 | 0.46 | 0.43 | **0.44** | 0.41 | 0.14 | 0.21 |
| 11056 | 0.70 | 0.57 | **0.63** | 0.62 | 0.59 | 0.60 | 0.68 | 0.37 | 0.48 |

To evaluate the practical impact, we performed 3D reconstructions using particles selected by each method (excluding EMPIAR-11056). On the remaining six datasets, the resolutions of maps from ParSeek particles were comparable to those from Topaz and CryoSegNet, with differences ≤0.5 Å from the CryoPPP reference on three datasets and ≤0.3 Å from Topaz on two others (Supplementary Table S3). Consistently, cross-correlation coefficients (Supplementary Table S4) showed that ParSeek achieved the highest values on EMPIAR-10028, 10093, 10345, and 10532, tying with CryoSegNet on 10028 (0.995) and outperforming both methods on the other three (0.941, 0.931, 0.920, respectively). Topaz performed best on EMPIAR-10017 (0.995), while CryoSegNet showed a slight advantage on the two conformers of EMPIAR-10081 (0.964, 0.950). Notably, on the challenging, orientation-biased EMPIAR-10532 dataset, CryoSegNet failed to produce an interpretable map (Supplementary Fig. S5) and yielded a lower correlation coefficient (0.614) compared to Topaz (0.911) and ParSeek (0.920). The reproducibility of resolution estimates was confirmed by minimal variation across triplicate reconstructions (Supplementary Fig. S6). Together, these results validate the reliability of ParSeek-picked particles for high-resolution structure determination.

Moving beyond global resolution, we assessed map interpretability by fitting established atomic models into four representative density maps: a large ribosomal complex (EMPIAR-10028, *Plasmodium falciparum* 80S ribosome)^28^, a symmetric soluble enzyme (EMPIAR-10017, β-galactosidase)^29^, a membrane-embedded ion channel (EMPIAR-10093, NOMPC)^30^, and a ligand-binding membrane receptor (EMPIAR-10081, HCN1)^31^. Across these four cases, the density maps allowed reliable fitting of the corresponding models. Specifically, the ribosome map resolved the RNA nucleotides and tryptophan side chains (Fig. 4a); β-galactosidase showed well-defined secondary structures (Fig. 4b); NOMPC revealed clear densities for its ankyrin repeat regions and central pore (Fig. 4c); and the HCN1 reconstruction captured ligand-induced conformational changes, where the A-, B-, and C-helices in the cyclic nucleotide–binding domain (CNBD) underwent a discernible shift upon cAMP binding (Fig. 4d).

**Fig. 4.**
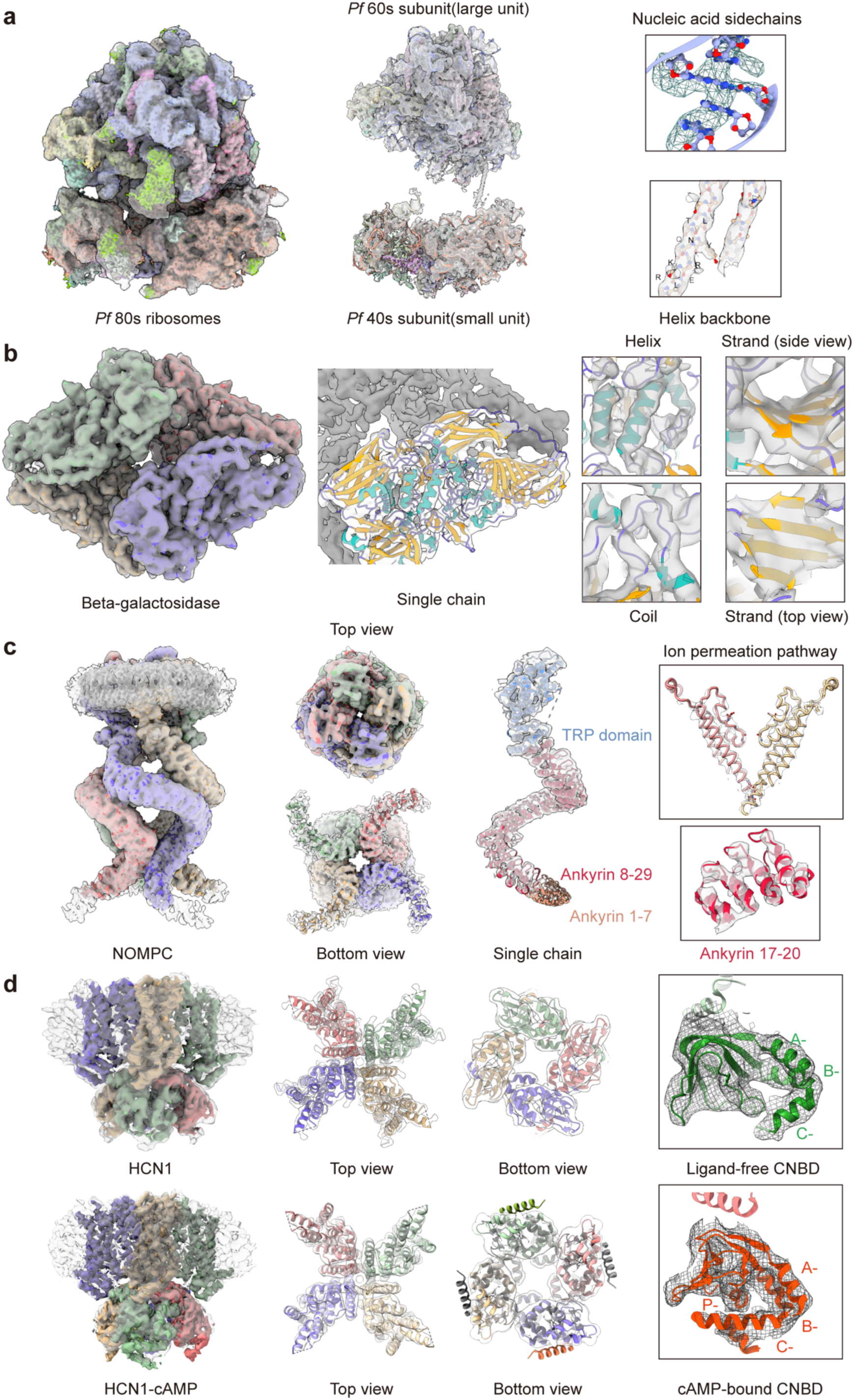
| Density maps reconstructed using particles picked by ParSeek and fitting of atomic models. **a**, Reconstruction of the *Plasmodium falciparum* 80S ribosome (EMPIAR-10028). Left, overall density map; Middle, atomic model fitted into the map (large subunit, PDB ID: 3J79; small subunit, PDB ID: 3J7A); Right, detailed view showing well-resolved density for representative RNA nucleotides and tryptophan side chains. **b**, Reconstruction of β-galactosidase (EMPIAR-10017). Left, overall density map. Middle, fitted atomic model (PDB ID: 3I3E) of one subunit within the symmetric oligomer; Right, detailed view highlighting well-defined secondary structure elements (α-helices, coils, and β-sheets). **c**, Reconstruction of the NOMPC mechanosensitive channel (EMPIAR-10093). Left, overall density map in side view; Middle left, top and bottom views of the map; Middle right, fitted atomic model (PDB ID: 5VKQ) of one subunit; Right, detailed view showing clear density around the ion permeation pathway and ankyrin repeats 17–20. **d**, Reconstruction of the HCN1 channel (EMPIAR-10081) in two functional states. Upper row, ligand-free state (PDB ID: 5U6O); Bottom row, cAMP-bound state (PDB ID: 5U6P). For each state: Left, overall density map in side view; Middle left, top view; Middle right, bottom view; Right, detailed view of the cyclic nucleotide–binding domain (CNBD) highlighting the conformational changes associated with cAMP binding.

These results demonstrate that ParSeek achieves competitive particle-picking accuracy, leads the F1-score on multiple benchmarks, and delivers reconstruction quality comparable to leading annotation-dependent methods. This shows that accurate cryo-EM particle picking can be achieved without manual labeling.

### Application validation of ParSeek using in-house cryo-EM experiments

Following quantitative benchmarks, we evaluated practical performance of ParSeek on our three experimental datasets: a reprocessed published dataset and two newly collected GPCR datasets. These datasets for high-resolution structure determination are: (i) a Tr67–Spike (BA.1) antigen–antibody complex from published data; (ii) an ICL3-BRIL-fused receptor CCR5 in complex with a proprietary ligand cpd1, for which we reported a new structure; and (iii) an LPC–GPR119–G_s_ complex at 2.8 Å resolution, which we revealed the ligand flexibility. All analyses used a single pre-trained model without dataset-specific tuning.

#### Cryo-EM structure of the Tr67–Spike complex

We first evaluated the ability to generalize across microscope configurations using a published dataset of Tr67–Spike complexes collected at 200 kV^32^. In this dataset, the computationally designed trivalent nanobody Tr67, which targets SARS-CoV-2 variants, is complexed with the Spike (BA.1) protein.

ParSeek processed 2,787 micrographs, and picked approximately 684,000 particles with high selectivity for ice holes and effective exclusion of carbon film regions (Supplementary Fig. S8a, b). The 2D classification yielded class averages representing multiple orientations, including a side view in which the Spike protein displays strong signal while the bound Tr67 nanobody is discernible but at lower contrast (Supplementary Fig. S8c). The 3D reconstruction achieved a resolution of 5.0 Å, an improvement over the previously reported structure (9.0 Å), and revealed that Tr67 binding stabilizes a three-RBD-up conformation of the Spike protein (Figs. 5a and Supplementary Fig. S8d). At this resolution, the map allowed domain-level interpretation: the N-terminal domain (NTD), receptor binding domain (RBD), and S2 subunit of Spike were unambiguously assigned, while weaker density permitted tentative placement of the Tr67 nanobody (Fig. 5b–d). Furthermore, protruding densities at known glycosylation sites were consistent with prior structural data (PDB ID: 8CYA), enabling modeling of N-acetylglucosamine (NAG) and β-d-mannose (BAG) glycans (Fig. 5e)^33^. This case demonstrates ParSeek can effectively treat the cryo-EM data collected at 200 kV.

**Fig. 5.**
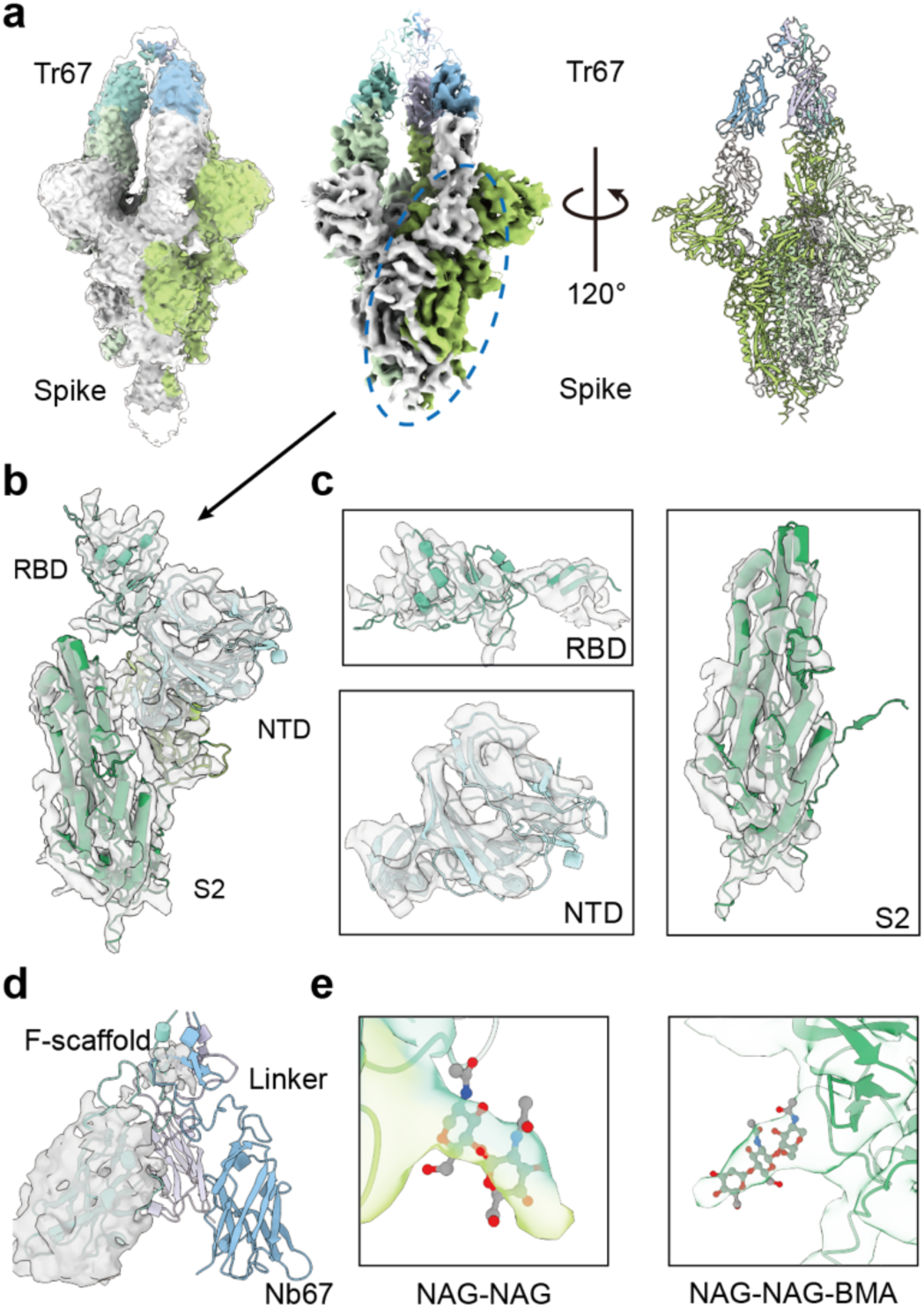
| Cryo-EM structure of the Tr67–Spike complex. **a**, Overall view of the 5.0 Å resolution cryo-EM reconstruction, showing the Spike complex in a three-RBD-up conformation bound by Tr67. **b**, Atomic model (helices as cylinders) of a single Spike protomer fitted into the corresponding cryo-EM density (gray surface), with the N-terminal domain (NTD), receptor-binding domain (RBD), and S2 domain indicated. **c**, Detailed views of the NTD, RBD, and S2 domains, showing the local cryo-EM density (gray surface) and fitted atomic model. **d**, Trivalent nanobody Tr67, with the atomic model for one chain superimposed on its cryo-EM map (gray surface). The Nb67 domain and scaffold show stronger density compared to the linker region. **e**, Detailed view of two glycosylation sites. The left site displays density consistent with an NAG-NAG disaccharide, while the right site shows density for an NAG-NAG-BMA trisaccharide.

#### Cryo-EM structure of cpd1–CCR5 complex

We next applied ParSeek to a dataset of the cpd1*–*bound CCR5 complex. For structural studies, CCR5 was stabilized by fusion with T4 lysozyme (T4L) at the N-terminus and with thermostabilized apocytochrome b562 (BRIL) in ICL3^34,35^. To further stabilize the complex and facilitate the cryo-EM data processing, BAG2, a BRIL-specific Fab fragment, was incorporated in complex purification^36^.

ParSeek picked 1.94 million particles from 4,450 micrographs. These particles were separated into two distinct populations in 3D classification: a major class which yielded a well-defined reconstruction at 4.0 Å resolution, enabling unambiguous modeling of most residues in the receptor, BRIL, and BAG2 (Figs. 6 and Supplementary Fig. S9); and a minor class that exhibited conformational variability, limiting high-resolution refinement (Supplementary Fig. S9a, c). This structural heterogeneity, along with the absence of density for the T4L domain in the major class, is likely attributable to the flexibility of the N-terminal fusion protein. Within the orthosteric binding pocket of the major class, we observed unambiguous density corresponding to cpd1. The density was consistent with a molecular scaffold derived from maraviroc, sharing its overall dimensions and volume (Fig. 6c)^34^. A comparative analysis with the CCR5 crystal structure (PDB ID: 4MBS) revealed an inward shift of about 8.0 Å at the intracellular end of helix VIII in the cpd1–CCR5 structure, as measured at the Cα atom of Q313^8^.^59^ (Fig. 6b)^37^. Compared to the apo–CCR5–G_i_ cryo-EM structure (PDB ID: 7F1S), the intracellular end of transmembrane helix 6 (TM6) in cpd1–CCR5 shifts inward by about 8.5 Å (measured at the Cα atom of R230^6.30^) due to the absence of G_i_ coupling (Fig. 6d). Meanwhile, helix VIII in our ligand-bound cpd1–CCR5 structure and the apo–CCR5–G_i_ complex adopt a similar conformation, which differs from that in the crystal structure. Thus, despite the inherent flexibility, ParSeek successfully identified functionally important structural details in this challenging membrane protein.

**Fig. 6.**
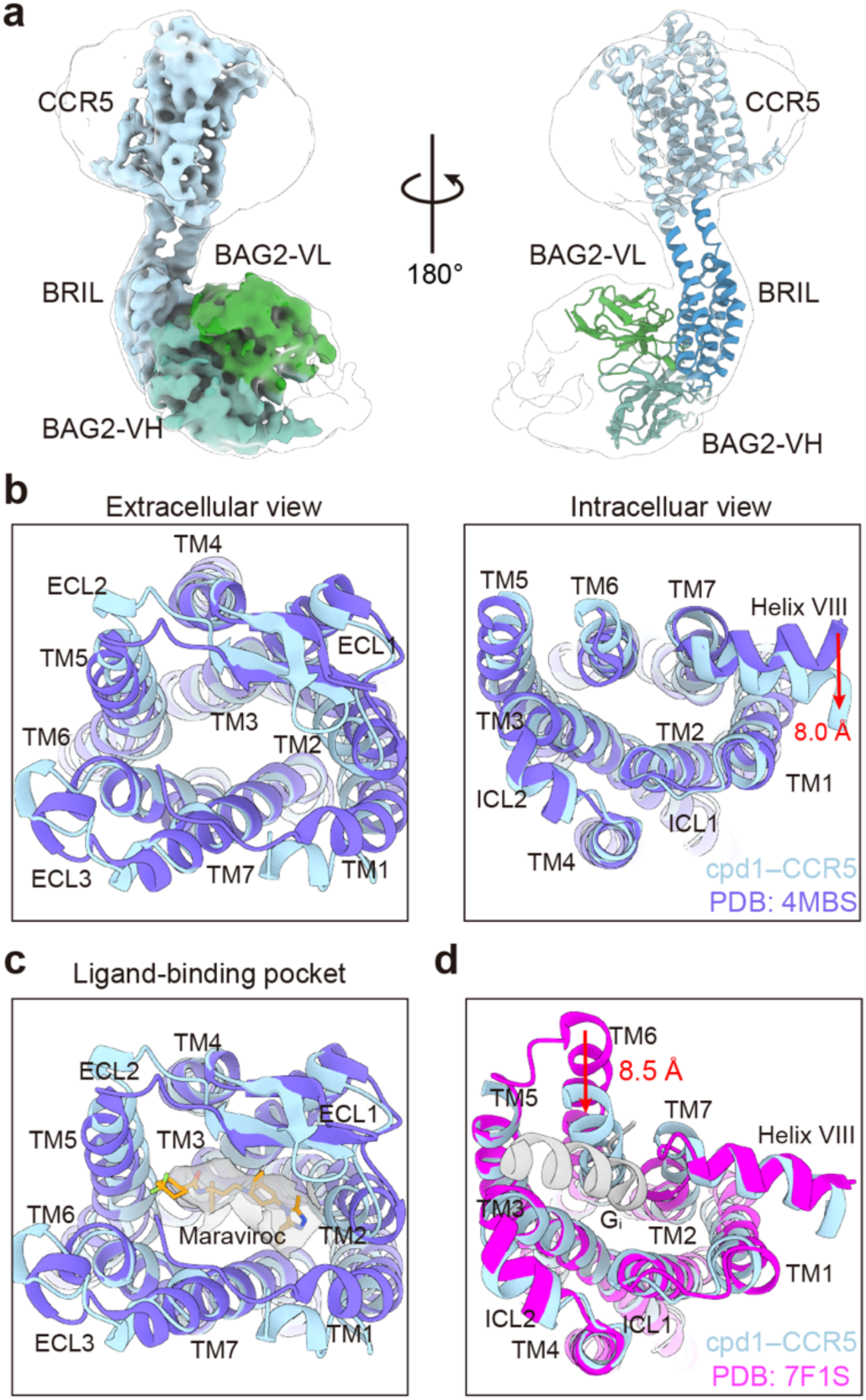
| Cryo-EM structure of the cpd1–CCR5 complex. **a,** Overall view of the 4.0 Å resolution cryo-EM reconstruction of the cpd1–CCR5 in complex with the BAG2 Fab fragment. The BAG2 Fab binds to the fusion protein BRIL. **b,** Structural superposition of the cpd1–CCR5 complex (blue) with the CCR5 crystal structure (PDB ID: 4MBS, purple), viewed from the extracellular (left) and intracellular (right) sides. The intracellular end of helix VIII in the cpd1–CCR5 structure exhibits an inward shift of approximately 8.0 Å. **c,** Close-up view of the ligand-binding pocket. The clear density map accommodates a proprietary drug candidate in a canonical binding pose, consistent with the inhibitor maraviroc (shown in stick representation). **d,** Structural comparison of cpd1–CCR5 (blue) with the apo-CCR5-G_i_ complex (PDB ID: 7F1S, magenta). The heterotrimeric G_i_ protein is shown in a gray cartoon. The intracellular end of TM6 in cpd1–CCR5 shifts inward by about 8.5 Å.

#### Cryo-EM structure of the LPC–GPR119–G_s_ complex

Finally, we evaluated ParSeek on a novel dataset of 5,643 micrographs featuring the LPC–GPR119–G_s_ complex. GPR119 is a class A GPCR and a potential therapeutic target for type 2 diabetes. The complex comprises the receptor bound to the engineered mini-G_si_ heterotrimer, which is a stable assembly consisting of a chimeric Gα subunit (with the N-terminus from Gα_i1_ fused to mini-Gα_s_), G_β1_ and G_γ2_, along with the stabilizing nanobody Nb35.

From 5,643 micrographs, ParSeek picked approximately 1.6 million particles. A subset of **∼**231,000 particles was selected for 3D reconstruction, yielding a 2.8 Å resolution structure that reveals the coupling interface between the activated GPR119 and the mini-G_s_ protein (Figs. 7, and Supplementary Fig. S10a). Notably, the map clearly shows density for the endogenous ligand lysophosphatidylcholine (LPC), thereby independently confirming the discovery by Xu et al.^38^, who reported a 3.1 Å resolution structure using a distinct dominant-negative G_s_ strategy (Fig. 7a,d). The high-quality map, with local resolution ranging from 2.2 to 4.0 Å (Supplementary Fig. S10d), allowed for detailed analysis of key structural features. First, we examined in the extracellular region, where the conserved disulfide bond between C155^45.50^ in TM3 and C78^3.25^ in ECL2 is visible with stronger cryo-EM density than in the reported structure (PDB ID: 7XZ5) (Fig. 7b), while four phenylalanine residues (F157^45.52^, F158, F161^5.35^ and F165^5.39^) form a hydrophobic extracellular region (Fig. 7c)^39^. Furthermore, the well-defined density for side chains across all transmembrane helices (TM1–TM7) resolves the core architecture of the receptor (Supplementary Fig. S10e). Notably, while our map confirms the overall active-state conformation, the absence of the density for the terminal of LPC acyl chain suggests increased conformational flexibility commencing from the carbonyl group, presenting a subtle difference in the detailed ligand geometry compared to the prior model (Figs. 7d and Supplementary Fig. S10e). This high-resolution density map, which slightly exceeds the previously reported GPR119–G_s_ structure, reveals fine structural details such as ligand flexibility and serves as evidence that ParSeek could deliver particle sets at a quality sufficient for near-atomic resolution refinement.

**Fig. 7.**
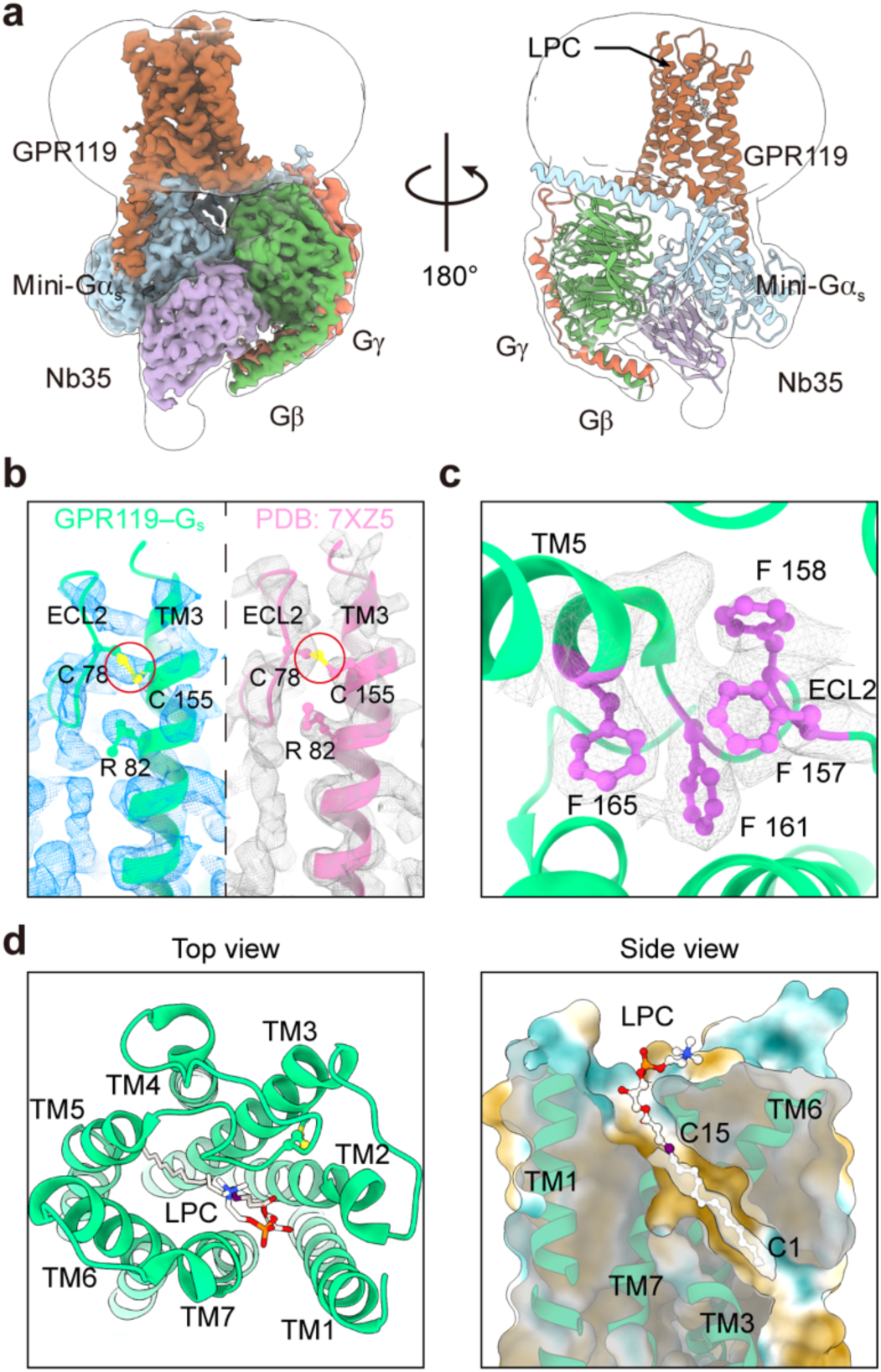
| Cryo-EM structure of the LPC*–*GPR119–G_s_ complex. **a**, Overall view of the cryo-EM density map (left) and the corresponding atomic model (right) of the LPC*–*GPR119*–*G_s_ complex. **b**, The cryo-EM density for the conserved disulfide bond (C155^45.50^–C78^3.25^, yellow sticks in red circles) is more defined in our GPR119–Gs structure (left) than in PDB ID: 7XZ5 (right). **c**, Extracellular hydrophobic region, primarily formed by ECL2 and enriched with phenylalanine residues (F157^45.52^, F158, F161^5.35^, and F165^5.39^; purple sticks). The cryo-EM density map (gray mesh) clearly defines the side chain densities. **d**, Orthosteric binding pocket of LPC. Top view (left) and side view (right) show clear density for the LPC molecule (sticks). The grey surface represents the well-defined density for the C1–C15 atoms (purple ball) of the LPC acyl chain, while the density for the remainder of the acyl chain is less well-defined.

## Discussion

Developing an accurate, automated particle picker has been hindered by the high cost of manual annotation and the biases inherent in reconstruction-based labeling. Here we present ParSeek, a deep learning picker trained entirely on synthetic data without any human annotation. By generating large-scale, objectively annotated training sets, ParSeek bypasses these obstacles. Our ablation experiments confirmed that using real buffer backgrounds and contamination annotations is critical for model robustness, while the projection-based ground truth avoids the selection biases inherent in manual or reconstruction-based labeling. ParSeek performs competitively with state-of-the-art methods and enables high-resolution structure determination from experimental data.

Previous strategies to reduce annotation burden range from semi-automated labeling (e.g., CryoPPP, CASSPER) to synthetic data as a supplement (Warp) or as a proof-of-concept (PARSED). More recently, realistic background generation (VirtualIce) improved synthetic image quality. These approaches are widely adopted because they are practical, computationally efficient, and often produce high-quality training sets for homogeneous samples. While these methods have advanced the field, each still depends partially on real annotations, custom heuristics, or simulation conditions that may not generalize. ParSeek differs by training exclusively on synthetic data derived from known 3D structures and experimental buffer backgrounds. ParSeek is not intended to replace existing pickers that rely on real labels, but rather to complement them where manual annotation is challenging, for example, in high-throughput screening, structurally heterogeneous samples, or initial model building for novel macromolecules. Moreover, ParSeek provides a standardized, bias-free benchmark for algorithm evaluation, as its training set is reproducible and independent of dataset-specific annotation choices.

Nonetheless, our approach also has limitations. ParSeek relies on known 3D density maps (e.g., from EMDB or PDB) to produce synthetic particles; for completely novel macromolecules without any homologous structures, the current pipeline cannot generate training data. Generating the synthetic dataset (6,600 micrographs) requires approximately 10 hours of GPU computation and about 250 GB of storage. Although this is an additional cost, it is far less labor-intensive than manual annotation and, as a one-time expense, the dataset can be reused for training or benchmarking.

Additionally, we validated ParSeek on complexes larger than 100 kDa, but we have not tested its performance on small (<40 kDa) or highly flexible proteins. Like most deep learning-based pickers, ParSeek currently uses downsampled micrographs (typically 4×) to balance computational efficiency and picking accuracy. However, this discards high-frequency information. Recent studies suggest that such high-frequency details may be critical for the reliable detection of small particles^40^, as they are required for successful reconstruction and alignment of sub-50 kDa complexes. Consequently, downsampling could potentially impair ParSeek’s ability to pick small targets, a limitation that cannot be compensated later. One future direction to address this is to reduce the binning factor (e.g., to 2× or 1:1) during both training and inference, thereby preserving high-frequency details. This would require re-training the model on full-resolution data and increase computational cost. Adaptive binning strategies might offer a practical trade-off.

Furthermore, while using realistic buffer backgrounds improves training fidelity, our current angular sampling is necessarily simplified for efficient synthetic data generation. For membrane proteins we biased sampling toward top and side views, but in reality particle orientation distributions are shaped by complex factors such as grid material, detergent concentration, and adsorption at the air-water interface^27,41,42^. This heuristic may limit generalization when applied to datasets with exceptionally strong or unusual orientation bias. One future direction is to replace the hand-crafted sampling with a data-driven approach, for example by learning orientation distributions directly from experimental micrographs or building statistical models informed by physical factors. Such strategies could better capture real-world heterogeneity and further improve the robustness of the resulting picker. Moreover, expanding the background library^43,44^ and restoring missing density signals^45^ are also important for further enhancing synthetic data fidelity.

In summary, this study demonstrates that a fully synthetic data strategy can train a cryo-EM particle picker with performance comparable to real-data methods, while eliminating the need for manual annotation. We anticipate that this synthetic-data paradigm can be extended to other steps in the cryo-EM pipeline, such as particle classification or 2D/3D refinement, paving the way for more automated and scalable structural biology workflows.

## Methods

### Generation of synthetic micrographs

In total, 897 background micrographs were downloaded from the public dataset (EMPIAR-12287)^18^. These micrographs were annotated with cryoSPARC Junk Detector and thus the contamination, crystalline ice, and carbon film regions were labeled (Supplementary Fig. S1). Synthetic 2D particle images were generated using 22 cryo-EM density maps, as listed in Supplementary Table S1. For three maps (EMDB-24900, 27752 and 32299) from CryoPPP that exhibited transmembrane signal loss, we substituted them with the maps (EMDB-31294, 38776 and 48592) featuring intact density (Supplementary Fig. S2). All the density maps were normalized using EMAN2^46^. Then, we applied different orientation sampling strategies based on protein type. For membrane proteins, structures were first oriented according to the OPM database^47^ to define their membrane plane. Subsequently, orientations were sampled as follows: 40% perpendicular to this plane (top/bottom views), 40% parallel to it (side views), and 20% via uniform spherical sampling. In contrast, for non-membrane proteins, orientations were generated entirely (100%) via uniform spherical sampling. Each map yielded a total of 1,692 unique orientations under its respective strategy.

Each density map was then rotated according to the sampled orientations, and a 2D projection was generated by integrating along the viewing direction. Each projection was then converted into a simulated particle image by applying the CTF corresponding to a randomly selected background micrograph and adding Gaussian noise (standard deviation estimated from background regions).

To assemble each synthetic micrograph, a background micrograph and a set of simulated particles images were first randomly selected. Particle images were then placed at random locations outside the pre-annotated contaminated regions. Finally, the contrast of the final composite image was adjusted based on the estimated ice thickness of the background micrograph.

For each synthetic micrograph, a corresponding ground-truth segmentation mask was generated. We first created a pure black image that matched the dimensions of the simulated micrograph. Then, a white circle was drawn at the coordinates of each valid particle placement. The diameter of each circle matched the known diameter of the corresponding particle (Supplementary Table S1).

For each of the 22 density maps, 300 synthetic micrographs were generated by randomly selecting background micrographs from the pool of 897. Each synthetic micrograph was accompanied by a ground-truth segmentation mask, resulting in a total of 6,600 unique micrograph–mask pairs.

### Neural network architecture and training of ParSeek

The ParSeek model was built upon a U-Net architecture integrated with a gated-attention mechanism (Supplementary Fig. S3a, b) for direct particle segmentation from micrographs^48,49^. The network consists of a 5-layer encoder, an intermediate layer, and a 5-layer decoder. In this design, the first encoder layer employs 11×11 convolution kernels to capture broad contextual features, while all subsequent layers use 3×3 kernels. A gated-attention module was incorporated in the final decoder layer to leverage morphological correlations among protein particles. This gated-attention module applies a sigmoid-activated weight map to the encoder features before concatenation, thereby performing a feature-wise selection (Supplementary Fig. S3b).

The model was trained on the synthetic micrographs described above using a composite loss function:

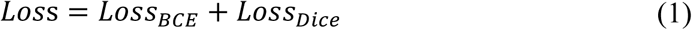

For the binary cross-entropy loss (𝐿𝑜𝑠𝑠*_BCE_*), a dynamic positive weight 𝜋 = |𝑏|/|𝑔| was utilized to address class imbalance between particle and background pixels. The training employed the AdamW optimizer with a learning rate of 0.001 and a weight decay of 0.01. For efficiency, input micrographs and masks were resized to 1024 × 1024 pixels and trained with a batch size of 8. The training process converged after 100 epochs on a single NVIDIA RTX 4070 Ti Super GPU, utilizing approximately 6 GB of memory (Supplementary Fig. S4a).

### Training of the control model (ParSeek-Noise)

A control model termed ParSeek-Noise was trained to evaluate the contribution of realistic background simulation. This model shared the same synthetic particle projections as the primary ParSeek model. To place these projections into micrographs, the particle coordinates were determined by KD-tree-based sampling with separation filtering, as described in the PARSED method^19^. Meanwhile, the background was simulated using additive Gaussian white noise at eight different SNR levels. A total of 6,600 synthetic micrographs were generated for ParSeek-Noise model, matching the scale of the primary ParSeek training set. The network architecture, training procedure, and all hyperparameters remained identical to those of ParSeek.

### One-frame self-supervised denoising model

A self-supervised blind-spot denoising network (BNN) based on the spatial-adaptive self-supervised (SASS) framework^48^ was employed to enhance the contrast of motion-corrected micrographs for subsequent particle picking. First, each raw micrograph was randomly cropped into 512 × 512 patches, which were normalized and augmented via rotation (0°, 90°, 180°, and 270°) (Supplementary Fig. S3c). These patches were then fed into a U-Net employing shiftConv2d layers^50^ to enforce the blind-spot condition (Supplementary Fig. S3d). Following the U-Net, the output underwent a post-processing step to produce the final denoised image 𝑋*_BNN_*. This involved padding, cropping, inverse rotation (to reverse the earlier augmentation), and a final convolutional layer (Supplementary Fig. S3e). The model was optimized using the L1 loss function and trained for 200 epochs with the Adam optimizer on a single NVIDIA RTX 4070 Ti Super GPU. The training used a learning rate of 0.001 and a batch size of 1 (Supplementary Fig. S4b).

### Benchmarking on CryoPPP datasets

Micrographs with corresponding ground truth particle coordinates from seven publicly available CryoPPP datasets were used to evaluate particle-picking performance. To evaluate ParSeek on these benchmark datasets, the following four-step processing pipeline was applied to each dataset: first, a self-supervised denoising model was trained and applied to all micrographs; second, the denoised outputs were processed by our segmentation model to produce probability maps, which were then binarized by applying a threshold of 0.1; third, the SAM model was employed to generate particle masks; and finally, particles were filtered by requiring a *predicted_iou* > 0.94 and a bounding box size of 90–110% of the expected particle diameter.

For comparative evaluation, CryoSegNet and Topaz were implemented according to their original publications and recommended workflows^7,11^. For CryoSegNet, micrographs were first preprocessed through Wiener filtering, non-local means denoising, CLAHE enhancement, and guided filtering. Denoised micrographs were then segmented by the author-provided U-Net, and particles were extracted as described in the original publication. For Topaz, micrographs were processed through normalization and 4× downscaling using topaz preprocess. Particles were then extracted with the pretrained ResNet16 model, with a radius of 12 pixels (15 pixels for EMPIAR-10017 and EMPIAR-10093) and a score threshold of 2.

To quantify picking performance, we calculated precision, recall, and F1-score. A predicted particle was considered a true positive if its center lay within a Euclidean distance of 0.3 times the particle diameter from any ground-truth particle center. To assess the quality of particles selected by each method, the corresponding particle set from each dataset was used for a separate 3D reconstruction. All reconstructions were carried out in cryoSPARC v4.7.0 using a consistent SPA workflow (a representative example for EMPIAR-10081 is shown in Supplementary Fig. S7). The resulting maps were assessed using Fourier Shell Correlation (FSC = 0.143) and local resolution estimation (with full results provided in Supplementary Figs. S5–6 and Supplementary Table S3). To further quantify the similarity between the reconstructed density maps and the cryoPPP reference map, we performed global fitting of the maps using emalign^51^, followed by local fitting with ChimeraX^52^, and finally computed the cross-correlation coefficients using Phenix ^53^(Supplementary Table S4).

### Protein expression

For structural studies of the cpd1–CCR5 complex, an engineered human CCR5 gene was cloned into a modified pFastBac1 vector. The construct includes an N-terminal HA signal peptide followed by a T4 lysozyme (T4L) fusion, and a C-terminal tag with a PreScission protease site, a FLAG tag, and 10× His tag. To stabilize the conformation, cytochrome b562RIL (BRIL) was inserted into the third intracellular loop (ICL3, between N226^6.26^ and E227^7.27^) by linkers derived from the A2A adenosine receptor (ARRQL at the N-terminus and ERARSTL at the C-terminus). Furthermore, a C-terminal truncation (ΔF320–L352) and four mutations (C58^1.60^Y, G163^4.60^N, A233^6.33^D, and K303^8.49^E) were designed to improve the thermostability and homogeneity of the sample. Recombinant baculovirus was generated using the Bac-to-Bac system and used to infect *Spodoptera frugiperda* (Sf9) cells at a density of 2 × 10^6^ cells/mL with an MOI of 5. Following a 48-hour incubation at 27 °C, the cells were harvested and stored at −80 ℃ for subsequent use.

For anti-BRIL Fab (BAG2) production, the corresponding gene was cloned into a pFastbac1 vector with an N-terminal GP64 signal sequence and a C-terminal 6×His tag. The construct was expressed in HighFive insect cells by infection at a density of 1.5 × 10^6^ cells/mL with an MOI of 5. After 48 hours of culture at 27 °C, the supernatant was collected by centrifugation.

To assemble the LPC-GPR119–G_s_ complex, a modified human GPR119 was engineered to include an N-terminal fusion comprising a hemagglutinin (HA) signal peptide and a thermally stabilized BRIL, followed by a C-terminal Flag tag and 2 × Strep tag. A point mutation (S237C) was introduced to enhance protein expression. For complex formation, a chimeric Gα subunit was constructed by replacing the N-terminal 18 amino acids of mini-Gα_s_ with the corresponding region from human Gα_i1_. Human G_β1_ and G_γ2_ subunits were also incorporated. All constructs were cloned into the pFastBac1 vector. GPR119, the chimeric Gα_i1_/mini-Gα_s_, G_β1_, and G_γ2_ were co-expressed in HighFive insect cells (Invitrogen). Cells were grown to a density of 1.5–2.0 × 10^6^ cells/mL and infected with baculovirus at a 5:3:3 MOI. Cells were harvested after 48 hours and stored at −80 ℃.

### Protein purification

#### cpd1–CCR5

The cell pellet from a 500 mL culture was thawed and resuspended in 50 mL of hypotonic buffer (10 mM HEPES pH 7.4, 20 mM KCl, 10 mM MgCl₂, protease inhibitor), followed by centrifugation at 14,000 × g for 30 minutes at 4 °C. The precipitate was washed with a high-osmolarity buffer (the hypotonic buffer supplemented with 1 M NaCl), and centrifuged at 14,000 × g for 30 minutes at 4 °C. The purified membrane was resuspended and solubilized for 3 hours at 4 °C in a buffer containing 25 mM HEPES pH 7.4, 150 mM NaCl, 0.375% (w/v) lauryl maltose neopentyl glycol (LMNG, Anatrace), 0.075% (w/v) cholesterol hemisuccinate (CHS, Sigma), 0.125% (w/v) glyco-diosgenin (GDN, Anatrace). After centrifugation (14,000 × g, 30 minutes), the supernatant was incubated with TALON cobalt-affinity resin (Takara) (overnight, 4 ℃). The resin was washed with a buffer composed of 25 mM HEPES pH 7.4, 150 mM NaCl, 0.0075% LMNG, 0.00075% CHS, 0.0025% GDN, and 20 mM imidazole. The protein was then eluted with 25 mM HEPES pH 7.4, 150 mM NaCl, 0.0075% LMNG, 0.00075% CHS, 0.0025% GDN, and 200 mM imidazole. Finally, the eluted protein was concentrated to 300 μL using a 100 kDa molecular weight cut-off (MWCO) filter.

#### BAG2

The supernatant was incubated with Ni-NTA resin (Clontech) resin for 1 hour at 4°C. The resin was washed with 10 column volumes (CV) of a buffer containing 20 mM HEPES pH 7.4, 150 mM NaCl, and 20 mM imidazole. The protein was then eluted with 5 CV of elution buffer (20 mM HEPES pH 7.4, 150 mM NaCl, 200 mM imidazole). The eluted Fab was concentrated and further purified by size-exclusion chromatography using a Superdex 200 Increase 10/300 column (GE Healthcare) pre-equilibrated with 20 mM HEPES pH 7.4, 150 mM NaCl. Finally, fractions were collected, concentrated to 5 mg/ml, flash-frozen in aliquots, and stored at -80°C.

#### Assembly of the cpd1–CCR5 complex

The purified cpd1–CCR5 was incubated with a two-fold molar excess of BAG2 on ice for 1h. The mixture was purified by size-exclusion chromatography on a Superdex 200 Increase 10/300 GL column (GE Healthcare) pre-equilibrated with 20 mM HEPES pH 7.5, 150 mM NaCl, 0.0075% LMNG, 0.00075% CHS, 0.0025% GDN to remove unbound components. The final complex was concentrated to 2.5 mg/mL using a 100 kDa MWCO concentrator.

#### LPC–GPR119–G_s_ complex

Cells were resuspended in lysis buffer (20 mM HEPES pH 7.4, 50 mM NaCl, 2 mM MgCl_2_, protease inhibitor), supplemented 10 μg/mL Nb35, and 25 mU/mL apyrase (1 hour, 20 ℃). After centrifugation (30,000 × g, 30 minutes), membrane proteins were solubilized in 25 mM HEPES pH 7.4, 150 mM NaCl, 0.5% LMNG, 0.025% CHS, 2 mM MgCl_2_, 25 mU/mL apyrase (2 hours, 4 ℃). Insoluble material was removed by centrifugation (30,000 × *g*, 30 min). The supernatant was incubated with Strep-Tactin resin (overnight, 4 ℃). The resin was washed with 20 column volumes of wash buffer (25 mM HEPES, pH 7.4, 150 mM NaCl, 0.01% LMNG, 0.0005% CHS, 2 mM MgCl_2_). The complex was eluted with elution buffer (200 mM Tris-HCl, pH 8.0, 150 mM NaCl, 0.01% LMNG, 0.0005% CHS, 2 mM MgCl_2_, 50 mM biotin) and concentrated using a 100 kDa molecular weight cutoff Amicon Ultra centrifugal filter. The concentrate was further purified by size-exclusion chromatography using a Superdex 200 Increase 10/300 column (GE Healthcare) pre-equilibrated with 25 mM HEPES pH 7.4, 150 mM NaCl, 0.01% LMNG, 0.0005% CHS, 2 mM MgCl_2_.

### Cryo-EM sample preparation and data collection

For the cpd1*–*CCR5 complex, 3 μL of sample at 2.5 mg/mL was applied to glow-discharged nickel–titanium (Ni-Ti) R1.2/1.3 300-mesh grids. Grids were blotted for 1 second at 100% humidity and 4 ℃ using a Vitrobot Mark IV (Thermo Fisher Scientific) and plunge-frozen in liquid ethane cooled by liquid nitrogen. Data were collected on a Titan Krios microscope (Thermo Fisher Scientific) operating at 300 kV, equipped with a Selectris energy filter (slit width 10 eV) and a Falcon 4 direct electron detector (Thermo Fisher Scientific). Movies were acquired in super-resolution counting mode using SerialEM. Detailed collection parameters are provided in Supplementary Table S5.

For the LPC*–*GPR119*–*G_s_ complex, the sample was concentrated to 3 mg/mL and 3 μL protein sample was applied to the glow-discharged holey grids (CryoMatrix R1.2/1.3, Au 300 mesh). The grids were bolted at 4 ℃ and 100% humidity for 1 second with a blot force of 0, and flash-frozen in liquid ethane cooled by liquid nitrogen using Vitrobot Mark IV. Data were acquired on a Titan Krios microscope operating at 300 kV, equipped with a BioQuantum energy filter (slit width 20 eV) and a K3 direct electron detector (Gatan), using EPU for automated data collection. Detailed parameters are summarized in Supplementary Table S5.

### Single-particle 3D reconstruction

Each of the three in-house cryo-EM datasets was independently processed through a standard SPA workflow. Briefly, ParSeek-picked particles from each dataset underwent 2D classification, *ab initio* reconstruction, and heterogeneous refinement in cryoSPARC. The resulting best classes were further refined using standard techniques as needed, including per-particle CTF refinement and motion correction, to produce final maps of 5.0 Å (Tr67–Spike), 4.0 Å (cpd1-CCR5), and 2.8 Å (LPC-GPR119–G_s_). Detailed workflows are provided in Supplementary Figs. S8–10.

### Model fitting and analysis

#### Tr67–Spike complex

The initial atomic model was assembled by aligning the separately built Tr67 and BA.1 Omicron Spike models to the reference structure (PDB ID: 8CYA)^33^ in ChimeraX^52^. The Tr67 model was constructed as described^32^, and the BA.1 Omicron Spike structure was predicted with AlphaFold2^54^. Glycan ligands from the reference structure were retained in the final assembled model. The initial atomic model was flexibly fitted into the density using Coot^55^.

#### cpd1–CCR5 complex

The initial model was generated using AlphaFold2^54^, rigid-body docked into the map, and then flexibly fitted using Coot^55^.

#### LPC–GPR119–G_s_ complex

The initial model was assembled using the GPR119 structure from PDB ID: 7WCN, the mini-Gs structure predicted by AlphaFold2^54^, and the LPC ligand coordinates from PDB ID: 7XZ5. This assembled model was then rigid-body docked into the density map and then adjusted in Coot and refined with Phenix^53^.

## Data availability

The data that support this study are available from the corresponding authors upon reasonable request. The cryo-EM density maps have been deposited in the Electron Microscopy Data Bank (EMDB) under accession codes EMD-80359 (Tr67–Spike), EMD-XXXXX (cpd1*–*CCR5), and EMD-XXXXX (LPC–GPR119–G_s_).

## Code availability

The source codes for ParSeek are publicly available via GitHub. And the address link is https://github.com/Fudan-HQLab/ParSeek.

## Acknowledgements

This work was supported by the National Natural Science Foundation of China (31971377, 31671386), and Shanghai Science and Technology Development Foundation Grant (22QA1412000).

## Author Information

These authors contributed equally: Jiaqiang Qian, Yousheng Gong.

## Contributions

J.Q. and Q. H. conceived the project. J.Q. developed the deep-learning model and source code. Y.G. performed cryo-EM sample preparation and acquired cryo-EM data. F.L. and Y.M. helped express and prepare the protein samples. J.Q. and Y.G. performed cryo-EM data processing and analysis. G.G. helped analyze the source code. Q. H. supervised the project and Y.Z. supervised F.L., Y.H., and Y.G. And J.Q., Y.G., F.L., Y.Z. and Q.H. wrote the draft of the manuscript. J.Q. and Q.H. revised the manuscript. All authors contributed to the review and editing of the manuscript.

## Competing interests

The authors declare no competing interests.

**Correspondence and requests for materials** should be addressed to Q.H. and Y.Z.

